# Dynamics of burst synchronization induced by excitatory inputs on midbrain dopamine neurons

**DOI:** 10.1101/2023.12.28.573502

**Authors:** Meng-Jiao Chen

## Abstract

Dopamine (DA) signals play critical roles in reward-related behavior, decision making, and learning. Yet the mainstream notion that DA signals are encoded by the temporal dynamics of individual DA cell activity is increasingly contested with data supporting that DA signals prefer to be encoded by the spatial organization of DA neuron populations. However, how distributed and parallel excitatory afferent inputs simultaneously induce burst synchronization (BS) is unclear. Our previous work implies that the burst could presumably transition from an integrator to a resonator if the excitatory inputs increase further. Here the responses of networked DA neurons to different intensity of excitatory inputs are investigated. It is found that as NMDA conductance increases, the network will transition from resting state to burst asynchronization (BA) state and then to BS state, showing a bounded BA and BS region in the NMDA conductance space. Furthermore, it is found that as muscarinic receptors modulated *Ca*^2+^ dependent cationic (CAN) conductance increases, both boundaries between resting and BA, and between BA and BS gradually decrease. Phase plane analysis on DA reduced model unveils that the burst transition to a resonator underpins the changes in the network dynamics. Slow-fast dissection analysis on DA full model uncovers that the underlying mechanism of the roles and synergy of NMDA and muscarinic receptors in inducing the burst transition emerge from the enlargement of nonlinear positive feedback relationship between more *Ca*^2+^ influx provided by additional NMDA current and more *I*_*CAN*_ modulated by added muscarinic receptors. Moreover, the lag in DA volume transmission has no effect on excitatory inputs-elicited resonator BS except for requiring more excitatory inputs. These findings shed new lights on understanding the collective behavior of DA cells population regulated by the distributed excitatory inputs, and might provide a new perspective for understanding the abnormal DA release in pathological states.

**Author summary:** The importance of DA signals is beyond doubt, so their encoding mechanism has very important biological significance and draws widespread attention. Yet the mainstream notion that DA cells individual provide a uniform, broadly distributed signal is increasingly contested with data supporting both homogeneity across dopamine cell activity and diversity in DA signals in target regions. Our article proposes that diverse distributed and parallel excitatory inputs can not only regulate the temporal dynamics of individual DA cell activity, but also simultaneously and synergistically regulate the network dynamics of DA cell populations by changing the local dynamics of DA cells, namely the burst transition from integrators to resonators. According to our perspective, many data that are difficult to interpret by the notion of the DA neuron individual coding can be well explained, such as burst asynchronization coding DA ramping signals, the scale of burst synchronization coding the amplitude of phase DA release, inhibitory DA autoreceptors facilitating resonator burst synchronization by postinhibitory rebound, etc. This study aims to elucidate the working mechanism of the DA system in physiological states such as positive reinforcement, and then to provide a new research perspective and foundation for understanding the abnormal DA release in pathological states.

## Introduction

Through signaling (1) obtained reward, (2) reward prediction, and (3) reward prediction errors (RPE) [1] [2], the midbrain DA system is integral to mediating reward-based neural activity underlying adaptive behavior to ensure individual survival and racial reproduction [3] [4] [5] [6]. Nowadays mainstream perspective suggests that DA signals are encoded by temporally changes in the neural activity of DA neurons individual, for example, the tonic DA release is encoded by low frequency firing and the phasic DA release is encoded by burst firing. Yet the notion that DA cells individual provide a uniform, broadly distributed signal is increasingly contested [7] [8] [9] with data supporting both homogeneity across dopamine cell activity [10] and diversity in DA signals in target regions [11] [12]. Therefore, an alternative perspective are gradually prevailed that besides of temporally changes in neural activity of DA cells individual, DA signals prefer to be encoded by spatially-organization of DA neurons population [13]. However, BS mechanism remains unresolved.

In recent years, a spate of elegant anatomic studies has reinvestigated the input/output of midbrain DA neurons using newer tracing methods, strikingly finding that the diversity and apparent non-specificity of afferent inputs to the DA cells arise widely from across the brain [11] [14] [15]. Fortunately, these diverse and parallel afferent inputs can be uniformly classified into two categories: (1) a largely excitatory inputs “sampling” distributed, parallel activity that drives DA activity, particularly bursting and (2) an inhibitory inputs that “gates” DA cells as a function of integrative discriminative selection process. When excitatory inputs are complemented by selective disinhibition, facilitate changes in DA cell activity. As these arise and increase across a large percentage of DA cells, synchrony emerges facilitating a population based phasic DA release.

Collectively, distributed and parallel excitatory inputs are the advocate factor in BS induction. Experimentally, by activating excitatory inputs, such as NMDA and muscarinic receptors [16] [17], reward associated stimuli [18] and reward-driven learning [19] indeed generate phasic DA release. Knock-out muscarinic receptors in mice VTA DA neurons attenuate phasic DA release [20], leading to defects in conditioned reinforcement for a cue previously paired with food reward [21]. Blockade of NMDA current, by local application of AP5 (NMDA antagonist) into the rat VTA or selective genetic inactivation of NMDA receptors in mouse VTA DA neurons, attenuates phasic DA release [22], leading to defects in reward-related learning [23]. Collectively, these studies are elucidating *the roles of NMDA and muscarinic receptors on BS induction in DA neurons population*. However, the dynamic mechanisms underlying these phenomena are still unclear. And intra-VTA co-infusion of carbachol (muscarinic receptors agonist) with AP5 abolishes phasic DA releases recorded in the NAc; however, co-administration of carbachol and NMDA into the VTA restore DA signals [24]. Again, this study has strongly suggested that *muscarinic and NMDA receptors probably synergistically induce BS between DA neurons population*. However, far less is known about the dynamic mechanisms underlying this synergy.

Contrary to intuition, DA neurons are not obligate synchrony, and accumulating evidences favor emergent synchrony [13]. Between midbrain DA cells, although direct connections (gap junctions) may be less prevalent, functional connections widely exist: D2 autoreceptors are coupled to G-protein activated inwardly rectifying *K*^+^ (GIRK) channels to produce an inhibitory postsynaptic current upon activation by dopamine [25]. Experimentally, it is reported that these D2 GIRKs paradoxically facilitate phasic DA release [26] [27]. However, researches on network dynamics unveil that, inhibitory connections tend to induce anti-phase BS [28] or cluster BS [29] which are adverse to induce phasic DA release compared to in-phase BS we concerned.

Since inhibitory connections go against (in-phase) BS induction, the burst firing of DA neurons (local dynamics of nodes) is then taken into consideration. According to Izhikevich’s research [30] [31], the bursts can be divided into integrators and resonators, which are decided by the bifurcation of the resting state of the bursts. Resonators are easier to synchronize than integrators, so when the bursts transition from integrators to resonators, BS will be evoked. Using slow-fast dissection analysis on the bursts induced by excitatory inputs in detail [32], our recent study has detected that in two slow variables state plane composed by intracellualr *Ca*^2+^ concentration (denoted by *Ca* for simplification) and gate variable *z* of SK current, although the trajectories go across saddle-node (SN) bifurcation curve to enter firing state (Figs. 1A and 1B, red curves), similar to other studies [33] [34], bifurcation curves of the resting state undergo some interesting changes. On DA neurons with normal SK conductance, the bifurcation curve of the resting state only consists of SN bifurcation (Fig. 1A, blue line); and while on DA cells with lower SK conductance, the bifurcation curve of the resting state consists of SN bifurcation (Fig. 1B, blue line) and Andronov-Hopf (AH) bifurcation (Fig. 1B, green line), transitioning via Bogdanov-Takens (BT) bifurcation (Fig. 1B, magenta dot). These results imply that although the burst is still an integrator after SK conductance moderately decreases, a further decrease in SK conductance may lead to the burst transition to a resonator.

**Fig 1.**
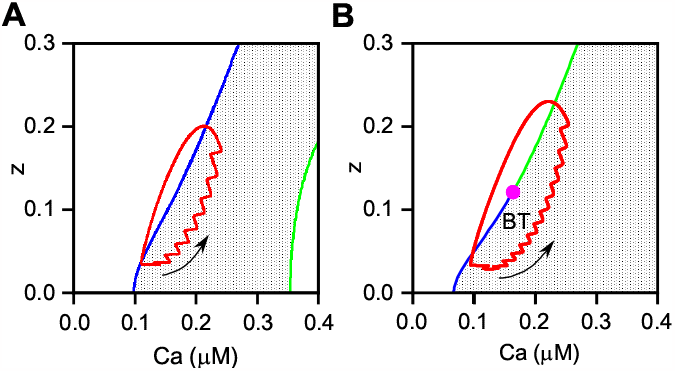
State-space trajectories (red curves) in *Ca*-*z* state space. Both trajectories enter firing state (grey zone) from resting state (white zone) through SN bifurcation curve. **(A)** On DA neurons with normal SK conductance there exists a SN curve (blue line) and a AH curve (green line), the former generates firing and the latter generates depolarization block. **(B)** On DA neurons with lower SK conductance there only exists a bifurcation curve of the resting state that generates firing which transit from SN (blue line) to AH (green line) bifurcation via BT bifurcation (magenta dot).

Molecular biology experiments have unveiled rapid translocation of TRPC channels into the cell surface after muscarinic receptors activation [35], resulting in CAN conductance increase [36] [37] which counteracts the effect of SK current, producing the phenomenon of a decrease in SK current [38]. Distributed cholinergic excitatory inputs from different brain region may act on the same DA neurons, thus affecting the activities of DA cells separately or simultaneously. When these cholinergic excitatory inputs are activated separately, moderate muscarinic receptors will be activated. And when they are activated simultaneously, adequate muscarinic receptors will be activated. Therefore, it is logically predicted that when adequate muscarinic receptors are activated, further increase in CAN conductance will occur, presumably producing a larger slow subthreshold depolarization, and then inducing more *Ca* accumulation which in turn will activate *I*_*CAN*_ further, that is, the positive feedback loop between *Ca* and *I*_*CAN*_ will be enlarged. Moreover, *z* of *I*_*SK*_ is *Ca* dependent and thus will increase with the increase of *Ca*, both together leading to state-space trajectory to shift up and right in *Ca*-*z* state space, ultimately resulting in the trajectory to enter the firing state through the AH bifurcation curve, namely the burst transition to a resonator.

Rapid *Ca*^2+^ chelation occludes NMDA-elicited phasic DA release [24], indicating that BS needs the necessary *Ca*^2+^ provided by NMDA receptors. Likewise, distributed glutamatergic excitatory inputs from different brain regions may also act on the same DA neurons, thus playing their roles separately or simultaneously. When glutamatergic excitatory inputs are activated simultaneously, adequate NMDA receptors will be activated. Therefore, it is easily predicted that more *Ca*^2+^ provided by adequate *I*_*NMDA*_ will open *I*_*CAN*_ further to produce a larger slow subthreshold depolarization, then in turn to induce more *Ca* accumulation, that is, the positive feedback loop between *Ca* and *I*_*CAN*_ will be enlarged once more. Additionally, *z* of *I*_*SK*_ will increase with the increase of *Ca*, both together resulting in the burst transition to a resonator.

Besides of above discussed same types of excitatory inputs, different types of excitatory inputs from different brain region can also simultaneously affect the activities of DA neurons [39]. Thus, it is naturally predicted that simultaneous activations of muscarinic and NMDA receptors will make it easier to enlarge the positive feedback loop between *Ca* and *I*_*CAN*_ than activation of one or the other receptors. Therefore, that BS will be induced as two receptors excited simultaneously whereas it will not happen as two receptors excited individually, should be happened.

To verify all predictions above, in the present study, a network composed of two DA neurons are constructed, which are coupled by functional inhibitory connection implemented by D2 GIRKs current. Each DA neuron contains a NMDA current which provides *Ca*^2+^ to calcium dynamics and a CAN current which is modulated by muscarinic receptors. Before NMDA and muscarinic receptors activation, the whole network will settle to the resting state after a short transient. In the following, to study DA neurons population responding to different intensity of same types of excitatory inputs, the electrical behaviors of coupled DA neurons are modeled by setting the maximal conductances of NMDA(CAN) current, 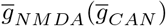, at moderate and adequate levels, respectively. To study DA neurons population responding to the synergistic effect of different types of excitatory inputs, moderate increases in 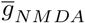 and 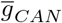 are done simultaneously. Then, the transition diagram of the network dynamics in the 2-D parameter space 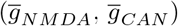 is analyzed. To uncover the underlying mechanisms of BS induction, phase plane analysis are conducted on the reduced isolated DA model. To unveil the underlying mechanism of the burst transition, slow-fast dissection analysis are done on the full uncoupled DA model. Finally, the effect of the lag of volume transmission on the BS induced by excitatory inputs is done.

## Materials and methods

### The Network Model

Our model of networked two neurons reads

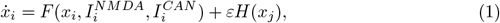

with *i, j* = 1, 2 the DA neuron (node) indices, *x*_*i*_ the state vector of the *i*th neuron, and *ε* the uniform coupling strength. 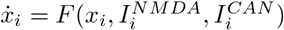 describes the local dynamics of the *i*th DA neuron, with *I*_*NMDA*_ and muscarinic receptor modulated *I*_*CAN*_. *H*(*x*) denotes the way how the neurons are coupled with each other, i.e., the coupling function. The DA cells are identical and the symmetrical inhibitory functional connection is fast and instantaneous. *ε* is dedicated by GIRK current *I*_*Kir*_ [40],

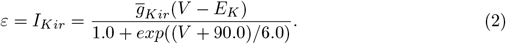

The parameter 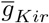 is the inhibitory coupling strengths and is set at 3*mS/cm*^2^, and the reversal potentials *E*_*K*_ = −90*mV* make the functional inhibitory. Recent study about DA transmission has revealed that although synaptic DA diffuses into extra-synaptic space, synaptic DA release is negligibly recognized by neighboring synapses [41]. Therefore, *H*(*x*) is modeled by the sigmoidal function,

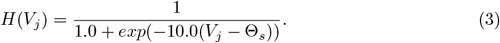

The coupling threshold Θ_*s*_ = −20*mV* is set to ensure that every spike in the single cell burst can reach the threshold. As a result, the spikes arriving from a presynaptic cell *j* activate the synapse current (through *H*(*V*_*j*_) switching from 0 to 1) entering the postsynaptic cell *i*.

Compared with synaptic transmission, DA volume transmission has relatively a longer lag after stimulation [42]. In order to study the effect of time lag on BS. Eq.(3) is changed to

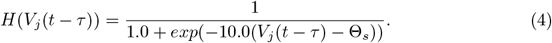

Here, *τ* is the time lag constant.

In simulations, our familiar isolated DA neuron dynamics is directly adopted from Ref. [32], which contains nine currents, in order of sodium, *I*_*Na*_(*V, h*), delayed-rectifier potassium, *I*_*DR*_(*V, n*), L-type voltage-gated calcium, *I*_*CaL*_(*V, dl, fl*), SK, *I*_*SK*_(*V, z*(*Ca*)), generic persistent potassium, *I*_*K*_(*V*), persistent sodium, *I*_*Nap*_(*V*), leak, *I*_*l*_(*V*), *Ca*^2+^-activated nonspecific, *I*_*CAN*_ (*V, Ca*), and NMDA, *I*_*NMDA*_(*V*), currents. Here, *V* represents the membrane potential of the *i*th DA cell; *h* is the inactivation variable for sodium current; *n* is the activation variable for potassium current; *dl* and *fl* are the activation and inactivation variables for L-type calcium current, respectively; *z* is the activation variable for SK current and is *Ca* dependent (see Ref. [32]); *Ca* is the intracellular *Ca*^2+^ concentration provided by *I*_*CaL*_ and *I*_*NMDA*_. Equations for gating variables and calcium dynamics used for simulation see Ref. [32] for details. The values of the parameters used for simulation are also directly adopted from Ref. [32] except some adjustments, all shown in Table 1.

**Table 1.** The values of the parameters.

| Parameter | Value | Parameter | Value | Parameter | Value | Parameter | Value |
| --- | --- | --- | --- | --- | --- | --- | --- |
| $\bar{g}_{Na}$ | 250.0 $mS/cm^2$ | $\bar{g}_{SK}$ | 3.5 $mS/cm^2$ | $E_{CAN}$ | 0 $mV$ | $C$ | 1 $\mu F$ |
| $\bar{g}_{KD}$ | 5.0 $mS/cm^2$ | $\bar{g}_{NMDA}$ | 0 $mS/cm^2$ | $E_{NMDA}$ | 0 $mV$ | $\tau$ | 100.0 $ms$ |
| $\bar{g}_K$ | 0.4 $mS/cm^2$ | $\bar{g}_{Kir}$ | 3 $mS/cm^2$ | $\varepsilon$ | $2.5 \times 10^{-3}$ | $C_{Mg}$ | 0.5 $\mu M$ |
| $\bar{g}_{NaP}$ | 0.002 $mS/cm^2$ | $E_{Na}$ | 55.0 $mV$ | $\kappa_1$ | 0.3 | $\Theta_s$ | -20 $mV$ |
| $\bar{g}_l$ | 0.015 $mS/cm^2$ | $E_K$ | -90 $mV$ | $\kappa_2$ | 2 | $\tau$ | 0 $ms$ |
| $\bar{g}_{CaL}$ | 0.075 $mS/cm^2$ | $E_{Ca}$ | 100.0 $mV$ | $\kappa_{SK}$ | 0.3 | | |
| $\bar{g}_{CAN}$ | 0.9 $mS/cm^2$ | $E_l$ | 0 $mV$ | $Ca_{basal}$ | 0.005 | | |

All parameters are directly adopted from Ref. [32], except shadowed ones which are adjusted, such as $\bar{g}_{SK}$ , $\bar{g}_{CaL}$ , $\bar{g}_{CAN}$ , and $\bar{g}_{NMDA}$ , and the rest of which are added, such as $\bar{g}_{Kir}$ , $\Theta_s$ , and $\tau$ .

### Phase plane analysis and fast-slow decomposition analysis

The system of isolated DA model is slow–fast such that the (*V, h, n, fl, dl*)-equations represent the 5-D fast “spiking” subsystem for the *i*th DA cell; the (*Ca, z*)-equations correspond to the slow 2-D “bursting” subsystem. *I*_*SK*_ gated by *z* and *I*_*CAN*_ controlled by *Ca* together produce slow subthreshold oscillation of *V*. When this slow subthreshold oscillation is above the threshold voltage (*V*_*th*_) of the fast subsystem, series AP generates, namely the burst emerges.

In order to do phase plane analysis to explore the underlying mechanism of BS induction, at the first, isolated DA neuron model is reduced into a 3-D model composed by (*V, Ca, z*) and the fast gating variables (*h, n, fl, dl*) are set to their corresponding steady states as a function of *V*. The reduced system is

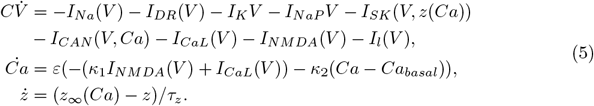

Then phase plane analysis on the reduced model under each cases are done. The position relationship between *Ca* nullclines and *V* nullclines at the *V*_*th*_ of fast subsystem determines the bifurcation of the resting state of the burst. That *Ca* nullclines is tangent to *V* nullclines, indicates the bifurcation of the resting state should be SN bifurcation, and thus the burst should belong to an integrator. That *Ca* nullclines intersects *V* nullclines, indicates the bifurcation of the resting state should be AH bifurcation, and thus the burst should belong to a resonator.

In order to unveil the underlying mechanism of the burst transition, fast-slow decomposition analysis on the full model under each case, by setting *z* as modification parameter and *Ca* as bifurcation parameter, are done. Then the bifurcation curves of the resting state and the state-space trajectories in *Ca*-*z* state space under different cases are compared.

Simulation is performed using Microsoft Visual *C* + + and bifurcation analysis after fast-slow decomposition is performed using the software *MatCont*. For different DA neurons, the initial values of state parameters in Eq. (1) are set to different values. The differential equations are solved using the fourth order Runge–Kutta method in steps of 0.01 ms (All codes see S1 File).

## Results

### BS induction by NMDA and muscarinic receptors, respectively

To investigate the effect of different intensity of NMDA receptors, co-activated by glutamatergic excitatory inputs from different brain regions, on the network behavior of coupled DA neurons, we start by exploring the variation of the network behavior with respect to the changes in 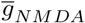, while keeping muscarinic receptor modulated CAN conductance fixed as 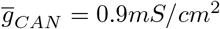. Setting 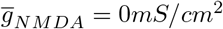, we plot in Fig. 2A the spatiotemporal evolution of the network after a transient period of *T* = 1.5 × 10^4^*ms*. It is seen that all the neurons are of steady membrane potentials, indicating that the state of network is completely resting. Increasing 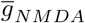 to 0.015*mS/cm*^2^, in Fig. 2B it is seen that all neurons turn to bursts but these bursts are not synchronous, i.e., the state of network transitions to BA. Documents indicate that BS induction requires a transient peak in glutamatergic excitatory inputs [43], this triggers our interest of looking for BS by increasing 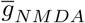 further. Fig. 2C shows the network evolution for 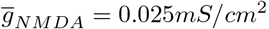. Here it is observed that all the neurons remain burst and these burst are synchronous, i.e., the state of network transitions to BS. As expected, BS is generated, and I use 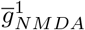and 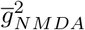to represent the transition boundaries from resting to BA and from BA to BS, respectively.

**Fig 2.**
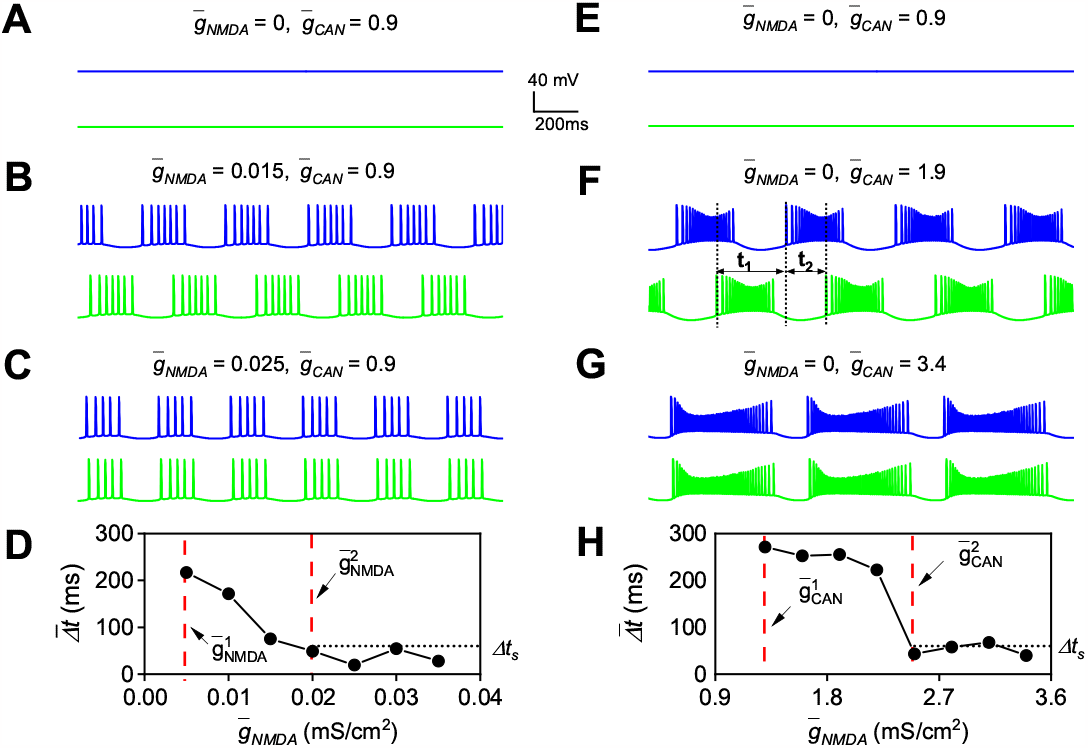
BS induced by NMDA (A-D) and muscarinic receptors(E-H), respectively. Fixing the CAN conductance as 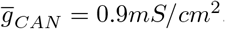, the spatiotemporal evolutions of the coupled DA neuron membrane voltages for 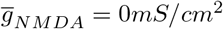 **(A)**, 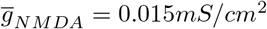 **(B)**, and 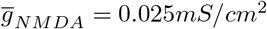 **(C). D** The variation of the average of minimum time difference between adjacent bursts of coupled DA neurons, 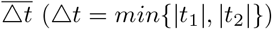, as a functions of 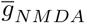. BA is evoked in the region 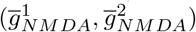, with 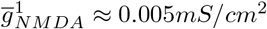 and 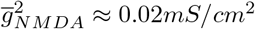, and because of 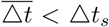 (Δ*t*_*s*_ is synchronization index and equals to 60 *ms*) BS occurs in the region 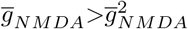. Fixing the conductance of NMDA receptors as 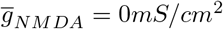, the spatiotemporal evolutions of the DA neuron membrane voltages for 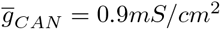 **(E)**, 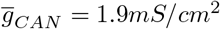 **(F)**, and 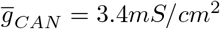 **(G). H** 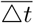 versus 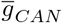. BA is happened in the region 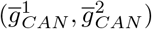, with 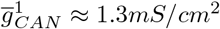 and 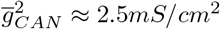, and BS occurred in the region 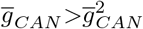. Blue and green represent cell 1 and cell 2, respectively.

To have more details on the transition of the network dynamics with respect to 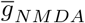, we plot in Fig. 2D the variation of the average of the minimum time difference between adjacent bursts of coupled DA neurons, 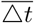, as a functions of 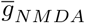. As shown in Fig. 2D, when 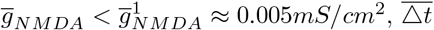 does not exist because the burst has not been induced yet; and then, during the interval 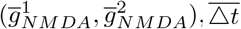 appears, and the value of 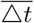 is gradually decreased from 210*ms* to below Δ*t*_*s*_ (synchronization index, Δ*t*_*s*_ = 60*ms*) at 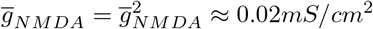; after that, 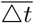 fluctuates below Δ*t*_*s*_. The transition scenario of the network dynamics therefore can be described as: resting → BA → BS, where the BA region of the network is identified as 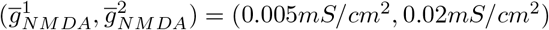, and the BS region of the network is identified as 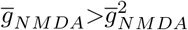. Figs. 2A-D clearly show the dependence of network dynamics on the intensity of NMDA conductance, indicating that the glutamatergic excitatory inputs from different brain regions can only induce the transition in DA neuronal network dynamics from BA to BS by simultaneously activating sufficient NMDA receptors.

To investigate the effect of different amount of muscarinic receptors, co-activated by cholinergic excitatory inputs from different brain regions, on the network behaviors of coupled DA neurons, we continue to explore the influence of the changes in CAN conductance modulated by muscarinic receptors, 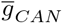, on network dynamics. Fixing the NMDA conductance at 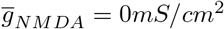, we plot in Figs. 2E-H the network evolution under different CAN conductance. When the CAN conductance is at initial level 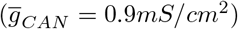, Fig. 2E shows that the network is at resting. Increasing 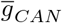 to 1.9*mS/cm*^2^, Fig. 2F shows that the network transitions to BA. When 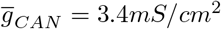, Fig. 2G shows that the network turns to BS. Again, we observe the transition from the resting to BA and then to BS by increasing 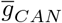. More details about the transition can be found in Fig. 2H, where 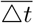 is plotted as a function of 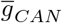, where it is shown that BA is induced in the region 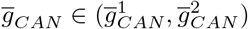, with 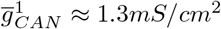 and 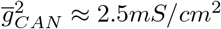; and BS is induced in the region 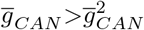. Figs. 2E-H show clearly the dependence of network dynamics on the amount of CAN conductance modulated by muscarinic receptors, indicating that cholinergic excitatory inputs from different brain regions can only induce the transition in DA neuronal network dynamics from BA to BS by simultaneously activating sufficient muscarinic receptors to modulate adequate CAN current.

### BS induction by NMDA and muscarinic receptors, synergistically

To verify that different types of excitatory inputs from different brain region can also simultaneously affect DA neuronal network dynamics, we directly adopt the cases showing in Fig. 2B and Fig. 2F. Clearly, when either moderate NMDA or muscarinic receptors are activated, only BA occurs. However, when moderate NMDA and muscarinic receptors are activated simultaneously 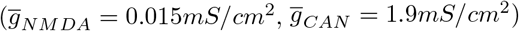, BS happens (Fig. 3). This result confirms that a synergistic effect exists between NMDA and muscarinic receptors in BS induction. Fig. 3 shows clearly the dependence of DA neuronal network dynamics on the synergy of NMDA and muscarinic receptors in BS induction, indicating that different types of excitatory inputs from different brain regions can also induce the transition of DA neuronal networks dynamic from BA to BS by simultaneously activating moderate NMDA and muscarinic receptors.

**Fig 3.**
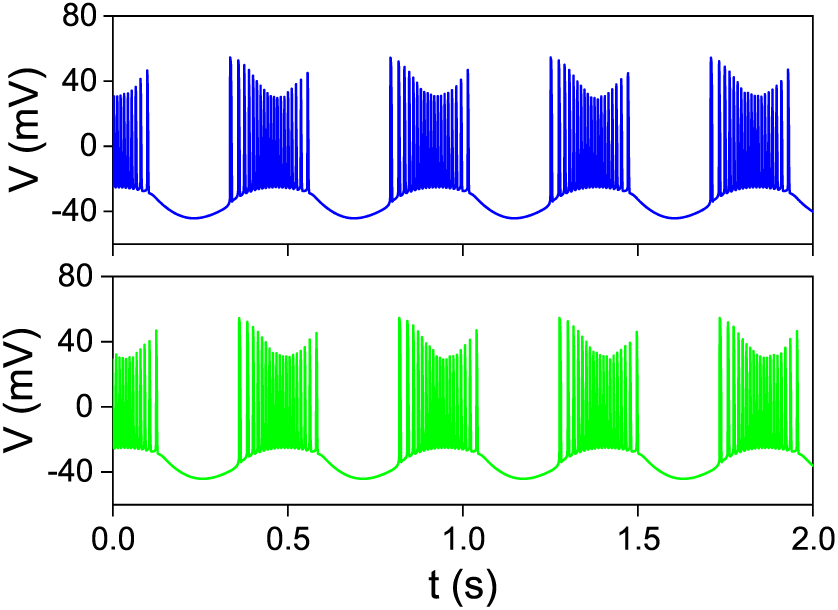
BS induced by NMDA and muscarinic receptors, synergistically. Blue and green represent cell 1 and cell 2, respectively.

### Bifurcation scenario on the 2-D parameter space 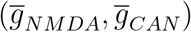

Results above show that same type of excitatory inputs with different intensities significantly induces different network dynamics, and different types of excitatory inputs can also induce different network dynamics simultaneously and separately. Therefore, it is necessary to globally describe the scenario of network dynamics in the 2-D parameter space 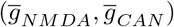. The results are presented in Fig. 4. For the sake of description, we directly adopt 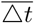 shown in Fig. 2D and 2H. Now, the influence of CAN conductance on the network behaviors can be systematically analyzed. It can be seen that as 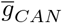 increases, the boundary separating BA state (Fig. 4, blue-green zone) from resting state (Fig. 4, white zone) and the boundary separating BS state (Fig. 4, red-orange zone) from BA state both gradually decrease.

**Fig 4.**
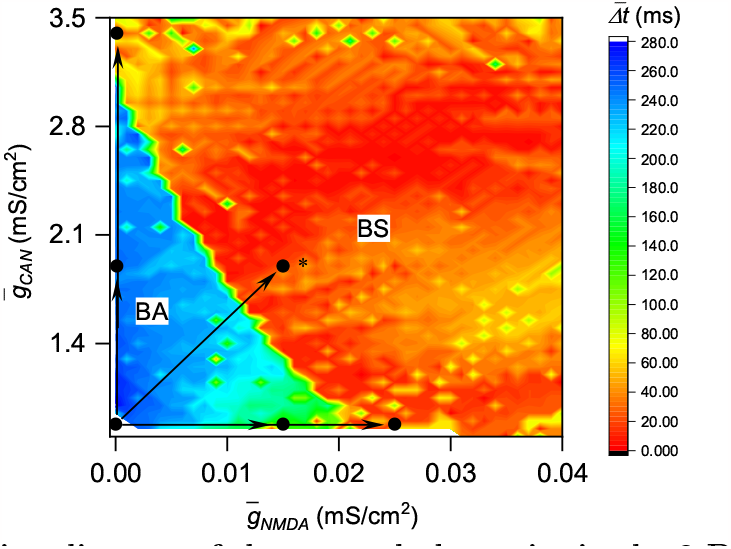
Bifurcation diagram of the network dynamics in the 2-D parameter space 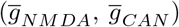. As 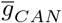 increases, the boundary from BS (BA) to BA (resting), 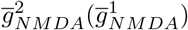, gradually shifts to left. White zone represents resting states, blue-green zone represents BA states, and red-orange zone represents BS states. Inserted six dots represent six cases of the network dynamics in control condition, during moderate and adequate NMDA(muscarinic) receptors activation, and moderate NMDA and muscarinic receptors simultaneous activation. The black horizontal (vertical) line with arrows denotes the transition process presented in Figs. 2**A-C** (Figs. 2**E-G**), and the black oblique line with arrow to the dot with ∗ denotes the transition process presented in Fig. 3.

As 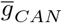 increases the boundary separating BA state from resting state gradually decreases, indicating that muscarinic receptors modulated CAN current is beneficial for NMDA-elicited BA induction. Our previous research has confirmed the synergistic effect of NMDA and muscarine receptors in the integrator burst induction, and its potential mechanism has also been revealed [32]. Similarly, as 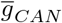 increases the boundary separating BS state from BA state gradually decreases, also indicating that muscarinic modulated CAN current contributes to NMDA-elicited BS induction. However, the potential mechanism of the synergistic effect of NMDA and muscarine receptors in inducing BS is not yet clear, and further investigation is needed.

## Mechanism analysis

So far, it is known that the form and strength of network connections between DA neurons have not changed, only external excitatory inputs that modulate the local dynamics of nodes have changed, illuminating us that the changes in the network dynamics of the DA neuronal network should be caused by changes in the local dynamics of the nodes, namely the burst should have undergone some kind of change. Therefore, in order to unveil the potential mechanism of BS induction, our focus is on analyzing the neuro-computational properties of the burst, especially the bifurcation of the resting state that determines whether the burst is an integrator or a resonator.

### Burst transition from an integrator to a resonator underpins BS induction

To analysis the bifurcation of the resting state of all bursts, phase plane analysis on the reduced DA model are done. Before phase plane analysis, the threshold voltage of fast subsystem, *V*_*th*_, is calculated. After removing slow currents *I*_*CAN*_ and *I*_*SK*_ and adding external DC current *I*_0_ to the fully DA neuron model, *V*_*th*_ is calculated out by adjusting *I*_0_. It is found that no matter how 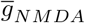 changes, *V*_*th*_ remains the same, approximate -38.7 *mV*.

Then, in each case, *z* is set as modification parameter, *Ca* nullclines and *V* nullclines are calculated, the results are shown in Fig. 5. Intuitively, changes in all cases are very similar, namely *V* nullclines shift downward as *z* decreases. When *z* stays at a higher level, *V* nullclines (Figs. 5, blue lines) intersects with *Ca* nullclines (Figs. 5, black lines) at two membrane potential levels located on both sides of *V*_*th*_ (Figs. 5, black dots), namely equilibria. The right one is unstable and the left one is stable, resulting in the entire system being in a resting state. When *z* decreases to a lower level, in the *V* space we concerned, *V* nullclines (Figs. 5, green lines) and *Ca* nullclines cannot contact each other, causing the entire system to be in a firing state.

**Fig 5.**
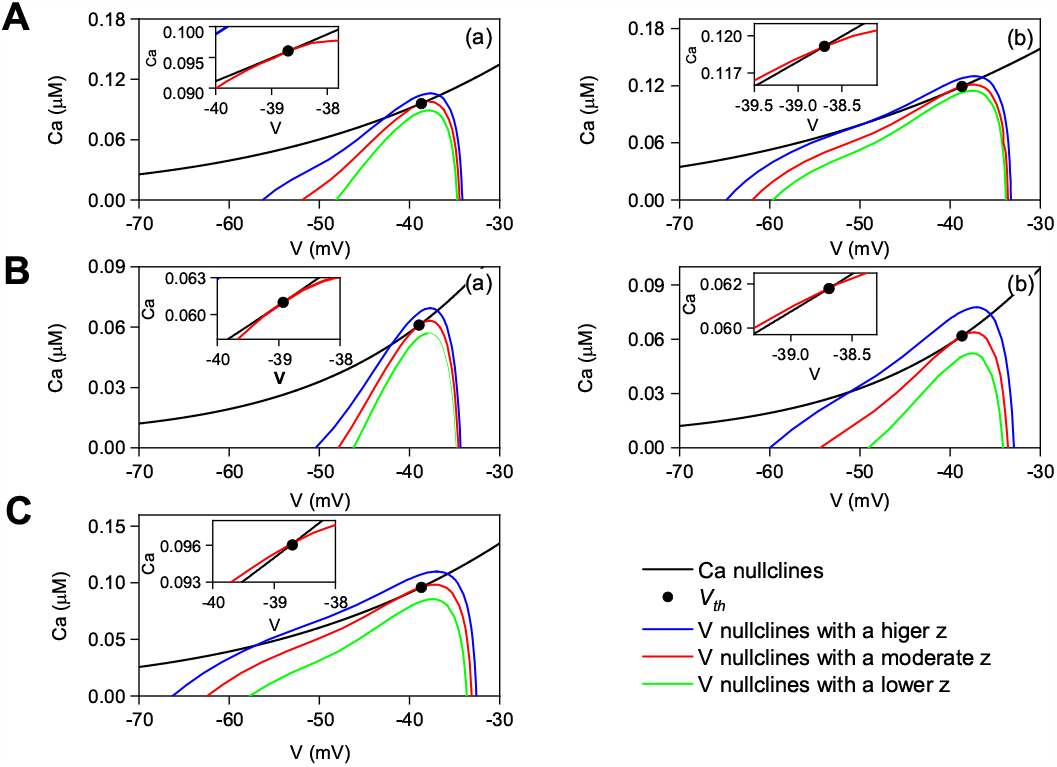
Phase plane analysis on each burst in reduced DA model. *Ca* nullclines (black lines) and *V* nullclines with a higher (blue lines), moderate (red lines) and lower (green lines) *z*, respectively, are shown for moderate **(a)** and adequate **(b)** NMDA receptors activation **(A)**; moderate **(a)** and adequate **(b)** muscarinic receptors activation **(B)**; and moderate NMDA and muscarinic receptors simultaneous activation **(C)**. Inserted graphs are expanded view near *V*_*th*_.

Then, a much closer look are taken to study the transitions from the resting state to the firing state, namely the bifurcation of the resting state. It is worth noting that around *V*_*th*_ the position relationship between *Ca* nullclines and *V* nullclines varies. In the cases of moderate NMDA or muscarinic receptors activation, *Ca* nullclines are tangent to *V* nullclines at *V*_*th*_ (Figs. 5 Aa and Ba, red lines); however, when adequate NMDA or muscarinic receptors are activated (Figs. 5 Ab and Bb, red lines), or moderate NMDA and muscarinic receptors are simultaneously activated (Fig. 5 C, red lines), *Ca* nullclines intersect with *V* nullclines at *V*_*th*_.

The tangent of *Ca* nullclines and *V* nullclines indicates that the bifurcation of the resting state is an SN bifurcation, confirming that the burst belongs to an integrator. The intersection of *Ca* nullclines and *V* nullclines means that the bifurcation of the resting state is an AH bifurcation, demonstrating that the burst is a resonator. Therefore, the results shown in Fig. 5 indicate that excitatory inputs generate BS by inducing the burst transition from an integrator to a resonator. Further research is needed on how excitatory inputs induce the burst transition.

### The enlargement of the positive feedback loop between *Ca* and *I*_*CAN*_ underpins the burst transition

The results of phase plane analysis indicate that excitatory inputs generate BS by inducing the burst transition. Therefore, in order to reveal the mechanism of the burst transition, slow subthreshold oscillations of those bursts and the results of fast-slow dissection analysis on those bursts are compared.

At the first, taken the burst induced by moderate NMDA receptors activation as an example to illustrate the process of fast-slow dissection analysis (Fig. 6A). With low initial *z* levels and *I*_*SK*_ deactivated, the resulting diagram shows that for each sufficiently low level of *Ca*, the stable feature is an equilibrium point, corresponding to the resting state. As *Ca* increases to a critical value *Ca*_*SN*_, an SN bifurcation occurs (Fig. 6Ba, blue dot), where a stable node coalesces with a unstable saddle points near - 38.6 *mV* and a new stable state corresponding to a periodic orbit emerges. However, as *z* gradually evolved and *I*_*SK*_ activated, an AH bifurcation occurs when *Ca* increases to a critical value *Ca*_*AH*_ (Fig. 6Bb, green dot). The coalescing of these stable and the corresponding unstable equilibrium points is illustrated in Fig. 6C, which superimposes the *V* -*Ca* bifurcation structures from Fig. 6B, showing that the stable branch on the left of *Ca*_*AH*_ expands to occupy a greater proportion of the *Ca* space (comparing curves a and b in Fig. 6C), and SN bifurcation transitions to AH bifurcation via BT bifurcation (Fig. 6C, magenta dot).

**Fig 6.**
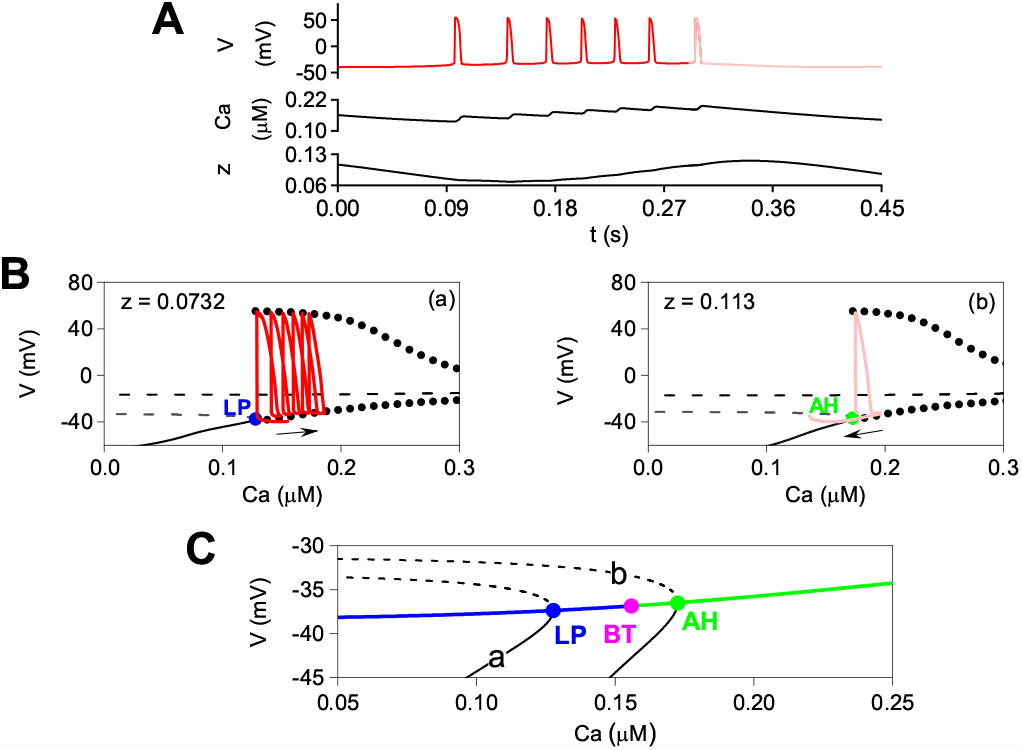
Bifurcation analysis on the burst induced by moderate NMDA receptors activation using fast-slow variables separation. **(A)** An expanded and color-coded burst voltage traces. **(B)** *V* -*Ca* bifurcation diagrams for fixed *z* values; red and light red trajectories correspond to **A**. Stable (unstable) equilibria are drawn with solid (dashed) lines, and periodic orbits (maximal and minimal voltages) are shown with dots. The branches at lower voltages that appear in the both *Left* **(a)** and *Right* **(b)** panels of **B**, consist of unstable (dashed lines) and stable (solid lines) equilibria, which intersect at a SN bifurcation (blue dot) and a AH bifurcation (green dot), respectively. **(C)** Expanded view near the SN and AH transitioning via BT bifurcation (magenta dot); **a** and **b** refer to *Left* **(a)** and *Right* **(b)** in **B**, superimposed without trajectories.

Then, I superimpose the *V* and *Ca* components of the solution trajectory onto the bifurcation diagram under different level of *z*. After the trajectory goes through the SN bifurcation with a lower *Ca*_*SN*_, entering periodic orbits, *Ca* level is high enough to maintain sufficient *I*_*CAN*_ activation and thus the positive feedback loop of *Ca* and *I*_*CAN*_ forms. Therefore, *Ca* continues to increase and the trajectory moves rightward in the *V* -*Ca* bifurcation diagram (Fig. 6Ba, red curves). However, the gating variable *z* evolves during the burst to evoke net outward current *I*_*SK*_ and the stable branch expands to occupy a greater proportion of the *Ca* space, thus the trajectory crosses AH bifurcation with a higher *Ca*_*AH*_ (Fig. 6Bb, light red curves) and the quiescent state becomes globally attracting, yielding burst termination, leading to a drop in *Ca* and to changing the direction of the trajectory along the *Ca* axis, and moving leftward relative to the *V* -*Ca* bifurcation diagram.

The quiescent state persists until *Ca*^2+^ clearance deactivates *z* of *I*_*SK*_, causing the stable branch to vanish via the SN bifurcation with a lower *Ca*_*SN*_, and the positive feedback loop of *Ca* and *I*_*CAN*_ forms, and thus the family of periodic orbits once again become globally attracting (a return to Fig. 6Ba). The resulting returned to low rate spiking initiates a new burst cycle.

After that, the bursts in the rest cases (Figs. 2C, 2F-G and Fig. 3) are analyzed by the same way. To be more intuitive, the dynamics of *Ca* and *z* of each burst in *Ca*-*z* plane are directly adopted to compare. Although all bursts and their subthreshold oscillations (Figs. 7 *Left*) exhibit different voltage trajectories, the dynamics of *Ca* and *z* of each bursts in *Ca*-*z* plane show some similar characteristics. In each case, in the *Ca* space we concerned, when fixed *z* is equal to *z*_*BT*_, BT bifurcation occurs (Figs. 7, magenta dots); for each fixed *z* less than *z*_*BT*_, the branch of quiescent states extends over a range of *Ca*, terminating when *Ca* reaches to the SN bifurcation curve (Figs. 7, blue lines); for each fixed *z* more than *z*_*BT*_, the branch of quiescent states expands to occupy a greater proportion of the *Ca* space, terminating when *Ca* reaches to the AH bifurcation curve (Figs. 7, green lines). In each case, during the burst, *z* exceeds 0, and thus the quiescent state exists for low *Ca*. However, the burst terminates only when *Ca* accumulates enough to activate *I*_*SK*_, which makes the trajectory to cross the AH bifurcation curve at a higher *Ca*_*AH*_ to enter the resting state, leading to a drop in *Ca* and change the direction of the trajectory along the *Ca* axis. Likewise, burst initiation occurs only when a decline in *z* of *I*_*SK*_ is enough and the positive feedback loop of *Ca* and *I*_*CAN*_ pushes the trajectory back across the bifurcation curve at a lower *Ca* to enter the firing state (Figs. 7, black and red cycles). Interestingly, under different cases, the bifurcation curves that the trajectories go across to enter the firing state are different.

**Fig 7.**
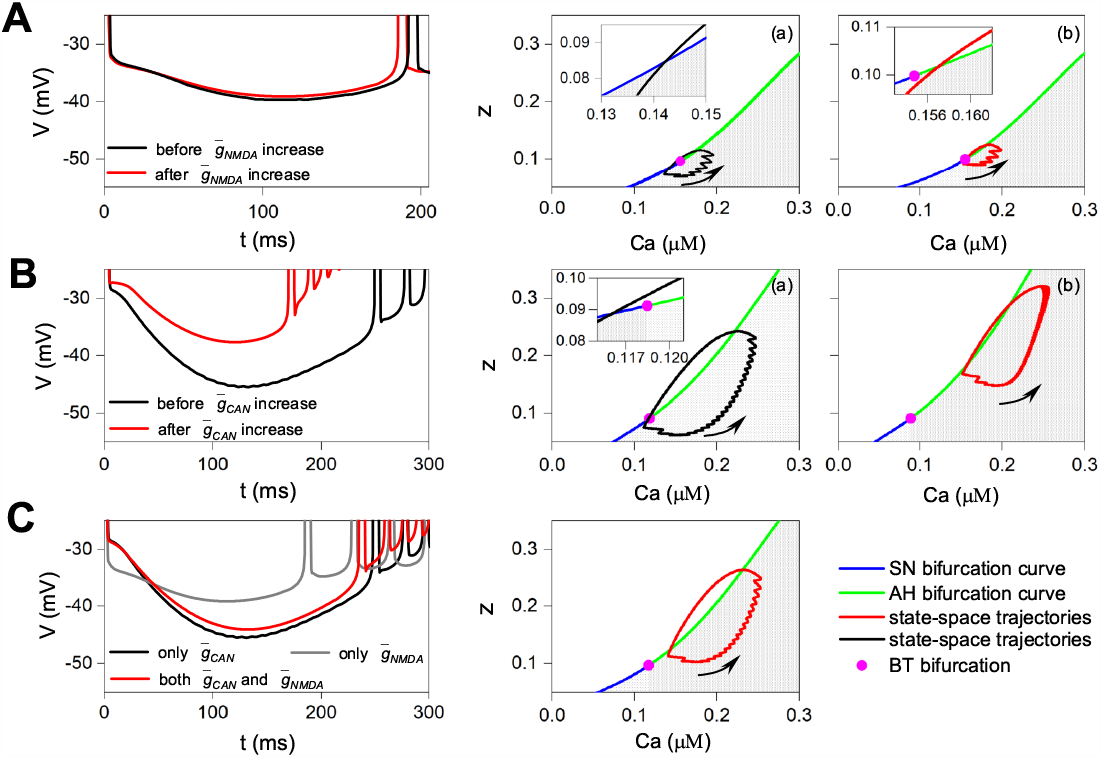
Mechanism analysis of the burst transition. Subthreshold voltage oscillations(*Left*) and state-space trajectories with SN, AH curves, and BT bifurcation are shown for moderate **(a)** and adequate **(b)** NMDA receptors activation **(A)**; moderate **(a)** and adequate **(b)** muscarinic receptors activation**(B)**; moderate NMDA and muscarinic receptors simultaneous activation **(C)**. In each state-space, white zone represents resting state and grey zone represents firing state. Inserted graphs are expanded view near BT bifurcation.

By comparison, it is found that when moderate NMDA receptors are activated, during the burst initiation, *z* declines to below *z*_*BT*_, the existing positive feedback loop of *Ca* and *I*_*CAN*_ pushes the trajectory back across the SN bifurcation curve (Fig. 7Aa, black cycle). However, when adequate NMDA receptors are activated, more *Ca*^2+^ is directly provided, which in turn further activates *I*_*CAN*_, resulting in a further depolarization of subthreshold oscillation (Fig. 7A, *Left*), and in turn further enhancing *Ca*; moreover, *z* is *Ca* dependent and thus increases with the increase of *Ca*, preventing *z* from declining to below *z*_*BT*_. Therefore, the enlargement of the positive feedback loop of *Ca* and *I*_*CAN*_ finally pushes the state-space trajectory up and right in *Ca*-*z* plane, thereby resulting in the trajectory to go across AH bifurcation curve to enter the firing state (Fig. 7Ab, red cycle), that is, the burst transition.

By same comparison, it is identified that when moderate muscarinic receptors are activated, during the burst initiation, *z* declines to below *z*_*BT*_, the existing positive feedback loop of *Ca* and *I*_*CAN*_ pushes the trajectory back across the SN bifurcation curve (Fig. 7Ba, black cycle). However, when adequate muscarinic receptors are activated, sufficient *I*_*CAN*_ directly induce a significant depolarization in subthreshold oscillation (Fig. 7B, *Left*), which in turn further activates *I*_*CaL*_ and *I*_*NMDA*_ to induce more *Ca* accumulation, and then in turn further activates *I*_*CAN*_; in addition, further increase in *Ca* once again prevents *z* from declining to below *z*_*BT*_. Therefore, the enlargement of the positive feedback loop of *Ca* and *I*_*CAN*_ finally leads to the burst transition (Fig. 7Bb, red cycle).

Above comparisons indicate that more increase in *Ca* provided by added NMDA receptors and more increase in *I*_*CAN*_ modulated by additional muscarinic receptors can induce the burst transition by enlarging the positive feedback loop of *Ca* and *I*_*CAN*_, respectively. These also suggest a synergy between *Ca* and *I*_*CAN*_ in enlarging the positive feedback loop, which should be the potential mechanism for synergistic induction of the burst transition by different types of excitatory inputs. To confirm this, I analyze the burst induced by moderate NMDA and muscarinic receptors simultaneous activations, and discover that it is indeed the case, that is, *Ca* provided by moderate NMDA receptors further activates *I*_*CAN*_ modulated by moderate muscarinic receptors, leading to a further depolarization of subthreshold oscillation (Fig. 7C, *Left*) and a further increase in *Ca*; the further increase in *Ca* once again prevents *z* from declining to below *z*_*BT*_. Therefore, the enlargement of the positive feedback loop of *Ca* and *I*_*CAN*_ results in the burst transition again (Fig. 7C, *Right*).

## Effect of the lag of DA volume transmission on BS

Although there is little discrepancy in transmission efficiency compared to synaptic transmission, the lag of DA volume transmission is relatively longer after stimulation (about 30-50ms) [42]. The effect of this lag on BS induced by excitatory synapses is still unclear. In order to study this effect, I directly adopt the new coupling function Eq. (4) which is added lag constant (*τ*) into Eq. (3). Then the effects of the time lag on the BS induced by NMDA or (and) muscarinic receptors are done by setting lag constant *τ* at different value.

Our simulation results indicate that BS induced by NMDA and muscarinic receptors exists all along as the value of lag constant *τ* gradually increases to 10*ms*, 30*ms*, and 50*ms*, respectively (Fig. 8A). Moreover, as *g*_*CAN*_ increases, the boundary between BS and BA still gradually decreases (Fig. 8A), demonstrating that the synergy between NMDA and muscarinic receptors in BS induction also exists all along. However, what has changed is that, as the lag constant *τ* increases, the boundary between BS and BA gradually moves to the right as a whole (Fig. 8B). These results together show that the lag of DA volume transmission has no effect on BS induction by NMDA or (and) muscarinic receptors; however, it makes BS induction more difficult, requiring more NMDA or (and) muscarinic receptors activation.

**Fig 8.**
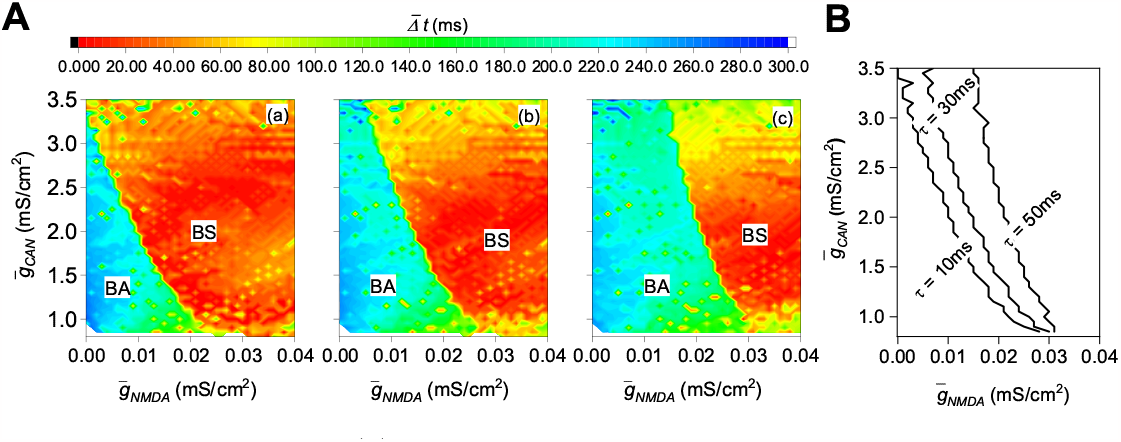
Effect of the lag (*τ*) of DA volume transmission on BS induced by excitatory inputs. **(A)** Bifurcation diagram of the network dynamics in the 2-D parameter space 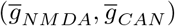 as *τ* = 10*ms* **(a)**, *τ* = 30*ms* **(b)**, and *τ* = 50*ms* **(c)**, respectively. **(B)** The boundaries between BS and BA under different lags.

## Discussions

A stark contrast is presented between midbrain DA input, highly diverse afferent inputs drawn broadly from across brain regions in which value related information does not appear to be neatly compartmentalized by region but rather widely distributed and intermixed, and the DA output system, which relies on relative uniformity of spiking activity across DA cell populations to transmit a simple scalar to target regions. So the transformation from heterogeneity of input to homogeneity of output is fundamental to DA’s function within the brain, and that the emergence of DA cell synchrony is the core mechanism mediating this function. Our research answers the questions of how heterogeneous inputs cause homogeneous outputs, i.e. distributed excitatory inputs induce BS, of DA neurons. Here, our study has found that when moderate excitatory receptors are activated, inducing integrator bursts that are difficult to synchronize, thus leading to BA; when adequate excitatory receptors are activated (regardless same types of excitatory receptors or different types of excitatory receptors), by enlarging the positive feedback loop between *Ca* and *I*_*CAN*_, inducing the bursts to transition from integrators to resonators that are easy to synchronize, thereby resulting in resonator BS. These results indicate that distributed excitatory inputs from different brain regions, whether of the same types or different types, cannot induce BS until they accumulate to a sufficient level in a large proportion of adjacent DA cells to induce the local dynamics of each DA neuron (node) to transition from an integrator to a resonator, thereby achieving the function of DA to convert heterogeneous inputs into homogeneous outputs.

In this study, BS is caused by changes in the local dynamics, that is, the bifurcation of the resting state of the burst transitions from SN bifurcation to AH bifurcation. In previous reports, for the burst, changes in local dynamics mainly focus on the bifurcation of limit cycle [44] [45] [46]. Actually, the transition in the bifurcation of the resting state has been observed on the spike firing [47], but related findings are rarely reported on the burst firing. Our research fills the gap that the transition in the bifurcation of the resting state can also happen on the burst. For the burst, transitions in two bifurcations of the resting state and limit cycle can both affect network dynamics, which should have rich, interesting, and even unexpected phenomena waiting for us to study.

Here, the burst transition is induced by the enlargement of the positive feedback loop between *Ca* and *I*_*CAN*_. Experimentally, somatodendritic [48] and axonal [49] synchronous DA release relies on large-scale high-intensity calcium oscillations and fast *Ca*^2+^ sensor synaptotagmin-1. Besides of *I*_*NMDA*_, calcium from other sources also contribute to *Ca* accumulation, such as voltage gated calcium channels (VGCCs) [50], nACh receptors [51], and intracellular calcium stores [52]. In addition to muscarinic receptors, other G-linked receptors can participate in regulating *I*_*CAN*_, such as mGluR1 [53] and CCK1 [54]. Moreover, besides of *I*_*CAN*_, an increase in other slow inward cation flux (such as *I*_*H*_ [43]) or a decrease in slow outward *K*^+^ flux (such as *I*_*SK*_ [55] and *I*_*M*_ [56]) can induce burst transition, and our data (see S1 Fig 1) has confirmed that a decrease in *I*_*SK*_ can induce this resonator BS in DA neuronal network. These all indicate that in addition to excitatory inputs, there are also other pathways to affect the resonator BS. Through these pathways, substances such as neuromodulators play roles in the DA system.

Experimentally, it is observed that inhibitory DA autoreceptors paradoxically facilitate phasic DA release [26] [27]. Theoretical studies have revealed that postinhibitory rebound promotes resonator spiking induction [31] and subsequently resonator spike synchronization [57] [58]. Naturally, it is speculated that DA autoreceptors should also promote the occurrence of resonator burst by postinhibitory rebound, thereby promoting resonator BS.

In our study, excitatory inputs always induce integrator burst before resonator burst, thus integrator BA always happens before resonator BS. Obviously, BS state triggers effective phasic DA release. So what is the significance of integrator BA’s existence? I have two speculations. One speculation is that, integrator BA would be the preparatory repository for resonator BS and the foundation for the scale change of resonator BS. Due to the prominent use-dependent depression, the burst of a single DA neuron cannot strongly enhance DA release [59] [60], and thus integrator BA plays little role in the generation of spatio-temporal effective DA phase release. Fig. 2 show that more glutamatergic (cholinergic) inputs induce resonator BS, and Fig. 3 show that cholinergic input facilitates glutamatergic inputs induced resonator BS, indicating that glutamatergic (cholinergic) inputs can regulate the scale of glutamatergic-(cholinergic-) induced resonator BS originating from other brain regions and cholinergic inputs can regulate the scale of glutamatergic-induced resonator BS, thereby producing different amplitude of phasic DA release. In experiments, it is observed that the amplitude of DA release varies due to different reward stimuli [61] [62]. The other speculation is that, BA state would induce DA ramping signals. Compared to the resting state (or low frequency tonic firing), although the burst of a single DA neuron cannot strongly enhance DA release, it still slightly enhances DA release. BA leads to this slightly increased DA releases asynchronous happen, resulting in DA gradual accumulation and then forming a DA ramping signal [63]. Additionally, because the integrator burst is more easily and generally induced, through some unclear mechanisms, if the integrator burst could directly synchronize (integrator BS) under the action of psychostimulants (e.g amphetamine), the scale of synchronization will be much greater than that of the resonator BS, then maybe leading to pathological DA release related to addiction.

## Conclusion

- Diverse external excitatory inputs from different brain regions cannot induce BS until they accumulate to a sufficient level in a large proportion of adjacent DA cells to induce the burst (local dynamics) of each DA neuron (node) to transition from an integrator to a resonator, thereby achieving the function of DA to integrate and convert heterogeneous inputs into homogeneous outputs.
- The roles and synergy of NMDA and muscarinic receptors (external excitatory inputs) in inducing the burst transition emerge from the enlargement of the nonlinear positive feedback relationship between more *Ca*^2+^ influx provided by additional NMDA current and more *I*_*CAN*_ modulated by added muscarinic receptors.
- The lag in DA volume transmission has no effect on excitatory inputs-elicited resonator BS except for requiring more excitatory inputs.

## Supporting information

**S1 Fig 1.**
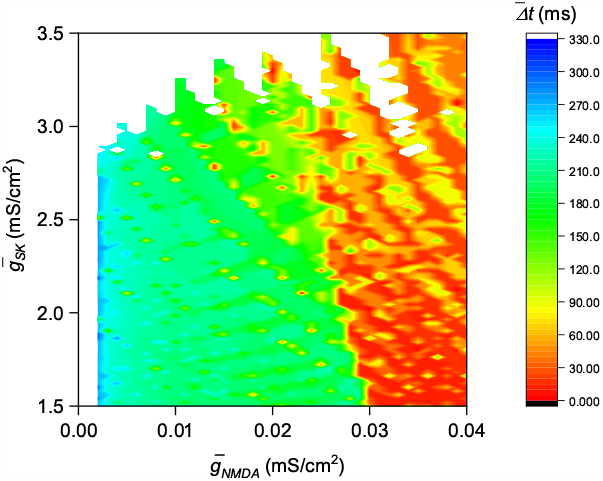
Bifurcation diagram of the network dynamics in the 2-D parameter space 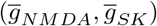.

**S1 File. All codes**.

## Acknowledgments

This work was supported by the Scientific Research Foundation of Key Laboratory of Sichuan Institute for Protecting Endangered Birds in the Southwest Mountains (BW202301), and by the Scientific Research Foundation of LeShan Normal University under the Grant Nos. RC202014 and JG2021-YB-04.

